# A biophysical and structural analysis of DNA binding by oligomeric hSSB1 (NABP2/OBFC2B)

**DOI:** 10.1101/2020.08.26.269084

**Authors:** Serene El-Kamand, Slobodan Jergic, Teegan Lawson, Ruvini Kariawasam, Derek J. Richard, Liza Cubeddu, Roland Gamsjaeger

## Abstract

The oxidative modification of DNA can result in the loss of genome integrity and must be repaired to maintain overall genomic stability. We have recently demonstrated that human single stranded DNA binding protein 1 (hSSB1/NABP2/OBFC2B) plays a crucial role in the removal of 8-oxo-7,8-dihydro- guanine (8-oxoG), the most common form of oxidative DNA damage. The ability of hSSB1 to form disulphide-bonded tetramers and higher oligomers in an oxidative environment is critical for this process. In this study, we have used nuclear magnetic resonance (NMR) spectroscopy and surface plasmon resonance (SPR) experiments to determine the molecular details of ssDNA binding by oligomeric hSSB1. We reveal that hSSB1 oligomers interact with single DNA strands containing damaged DNA bases; however, our data also show that oxidised bases are recognised in the same manner as undamaged DNA bases. We further demonstrate that oxidised hSSB1 interacts with ssDNA with a significantly higher affinity than its monomeric form confirming that oligomeric proteins such as tetramers can bind directly to ssDNA. NMR experiments provide evidence that oligomeric hSSB1 is able to bind longer ssDNA in both binding polarities using a distinct set of residues different to those of the related SSB from *Escherichia coli*.

## Introduction

Cells are subjected to constant oxidative stress that can damage the integrity of the genome and lead to diseases including cancer ^1,2^. To maintain genomic stability, oxidative DNA damage must be repaired before replication and cell division can occur.

The oxidation of guanine, the most commonly modified DNA base, results in the formation of the highly mutagenic 8-oxo-7,8-dihydro-guanine (8-oxoG) ^3–6^. Additional to the canonical Watson-Crick base pairing with cytosine, 8-oxoG has the ability to form stable Hoogsteen pairs with adenine, meaning that a G:C to A:T transversion may occur during replication as a result of this base modification ^7^.

Base excision repair (BER) is the main mechanism responsible for preventing the build-up of 8-oxoG in the human genome ^8^. In the BER process, human oxoguanine glycosylase 1 (hOGG1) first acts as a DNA glycosylase, cleaving the *N*-glycosidic bond, then functions as an apurinic/apyrimidinic (AP) nuclease, removing the 3’ phosphate of the resulting abasic site ^9^. DNA polymerase beta (POLβ) replaces the removed 8-oxoG nucleotide with a guanine residue that is ligated in place by the action of DNA ligase III ^10^.

Single stranded DNA binding proteins (SSBs) bind to and protect ssDNA in DNA processing events including DNA repair ^11,12^. Human SSB1 has previously been found to function in double strand break (DSB) repair, forming part of the sensor of single-stranded DNA complex 1 (SOSS1 complex) ^13–15^. More recent studies have found that hSSB1 is also involved in the removal of 8-oxoG from the genome, playing a central role in the recruitment of hOGG1 to damaged chromatin after oxidative damage ^16^.

hSSB1 monomers contain a single, highly conserved OB fold at the N-terminus followed by a flexible spacer region and a conserved C-terminal tail that is involved in protein-protein interactions ^17,18^. There are three cysteine residues present in the OB fold of hSSB1, two of which are solvent exposed (C81 and C99) ^19^. These cysteine residues, along with a set of charged and polar residues within the OB domain of hSSB1, facilitate the oligomerisation of hSSB1 under oxidative conditions, where hSSB1 forms homo-tetramers and larger oligomers ^20,21^. Our model of hSSB1 oligomerisation has also revealed that monomer-monomer and dimer-dimer interactions stabilising the hSSB1 tetramer occur at distinct surfaces of the OB domain that do not overlap with the ssDNA binding surface, indicating that ssDNA recognition by stable hSSB1 tetramers may be possible.

However, despite its significant functional importance, the exact molecular details of DNA binding by oligomeric hSSB1 remain unknown. In this study, we utilise solution-state nuclear magnetic resonance (NMR) spectroscopy and surface plasmon resonance (SPR) to elucidate the structural and biophysical details of oligomeric hSSB1-DNA interaction.

## Materials and Methods

### Protein expression and purification

Expression of GST-tagged hSSB1 (residues 1-123 encompassing the OB domain) was induced in *E. coli* BL21(DE3) by the addition of 0.2 mM IPTG for 15 h at 25°C ^20^. Isotopically labelled ^15^N-hSSB1_1-123_ was prepared in a fermenter to allow for nitrogen levels to be monitored and the source of nitrogen to be changed to ^15^NH_4_Cl before expression was induced. Cells were lysed by sonication in lysis buffer (10 mM MES, pH 6.0, 50 mM NaCl, 0.5 mM PMSF, 0.1% Triton X 100 (+ 3 mM TCEP for monomeric hSSB1)). Lysed cells were centrifuged, and the supernatant was subjected to GSH affinity chromatography followed by HRV-3C protease cleavage, leaving a five residue (GPLGS) non-native overhang at the N-terminus. Eluate from GSH affinity chromatography was subjected to heparin affinity chromatography using a HiTrap HP Heparin column (5 mL GE Healthcare) equilibrated with NMR buffer (10 mM MES, pH 6.0, 50 mM NaCl (+ 3 mM TCEP for monomeric hSSB1)). The hSSB1 protein was eluted from the column by increasing salt concentration (50-1000 mM NaCl gradient). Fractions correlating to a distinct peak in UV absorbance (280 nm) were collected and analysed by SDS-PAGE. For SPR, the protein was buffer exchanged into SPR buffer (50 mM Tris, pH 7.6, 50 mM NaCl, 0.4 mM EDTA, 0.005% P20 (+ 0.5 mM DTT for monomeric hSSB1)). Protein concentrations were determined using the absorbance at 280 nm and the theoretical molar extinction coefficient for hSSB1_1-123_ (11460 M^−1^cm^−1^). Note that all protein concentrations in this study refer to hSSB1 monomers, as oxidised hSSB1 consists of a mixture of monomers, dimers, tetramers and higher oligomers that cannot be separated ^21^.

### NMR spectroscopy

Purified protein samples (80-300 µM) were prepared in NMR buffer with 10% deuterium oxide (D_2_O) and 20 µM 4,4-dimethyl-4-silapentane-1-sulfonic acid (DSS). One dimensional (1D) and two dimensional (2D) spectra were acquired on a Bruker Avance 600 or 800 MHz spectrometer at 25°C using standard 1D and 2D pulse programs, respectively. All spectra were processed using TOPSPIN (Bruker, Biospin) with reference to the chemical shift of DSS (0 ppm) and assignments from our previous studies ^20,21^ were transferred using Sparky (T. D. Goddard and D.G. Kneller, University of California at San Francisco). The calculation of weighted chemical shift changes was carried out as described ^22^ using OriginPro 2019 (OriginLab Corporation, Northampton, MA, USA). All ssDNA oligomers were dissolved in NMR buffer prior to recording NMR experiments.

### Surface Plasmon Resonance (SPR)

All SPR experiments were carried out on a BIAcore T200, utilising the biotin-streptavidin system. Biotinylated oligo(dT)_12_ ssDNA with a 3A spacer [5′-btn-AAATTTTTTTTTTTT-3′] was injected into flow cell 2 of a streptavidin coated sensor (SA) chip at a flow rate of 5 µL/min for 190 s in SPR buffer. Approximately 88 response units (RU) of DNA was immobilised onto the surface. An unmodified flow cell 1 acted as a control. Solutions of purified hSSB1_1-123_ were injected for 20 s (monomer) or 1000 s (oligomer) at a flow rate of 20 µL/min. Flow cells were regenerated with 1 min injections of 1M MgCl_2_ (monomer) and 2 min injections of 2 M MgCl_2_ (oligomer) at 5 µL/min. Data obtained from SPR experiments were analysed using BIAevaluation 4.1.1. Regions chosen to determine equilibrium values were selected at the point of saturation. The dissociation constant (*K*_D_) and the theoretical response for binding of analyte at saturation to immobilised ligands (*R*_max_) of each interaction was derived from the correlation between the equilibrium values of the binding curve, and the hSSB1_1-123_ (monomer) concentration by fitting data to the Langmuir equation. All SPR figures were created using OriginPro 2019.

### Paramagnetic relaxation enhancement (PRE) labelling

Paramagnetic relaxation enhancement (PRE) NMR techniques using paramagnetic-labelled oligonucleotides (PRE-oligo(dT)_6_) [5′-G’(PRE)’GTTTTTT-3′] were utilised to determine the orientation in which ssDNA binds to hSSB1. Synthesis and purification of the PRE-labelled ssDNA was carried out in analogy to the study on the SSB from *Sulfolobus solfataricus* (SsoSSB) ^23^. However, in contrast to the described method ^23^, a reduced version of the paramagnetic-labelled oligonucleotide could not be used in this study due to the reducing agent impacting on the oligomeric state of the hSSB1 protein. Thus, an unlabelled oligonucleotide [5′-G’GTTTTTT-3′] was used as a control. Both PRE- and control ssDNAs were prepared in NMR buffer. The ratio of the peak intensities of hSSB1 bound to control DNA versus PRE-labelled DNA from HSQC spectra were calculated using Sparky (T. D. Goddard and D.G. Kneller, University of California at San Francisco) and plotted in OriginPro 2019 (OriginLab Corporation, Northampton, MA, USA).

## Results

### The presence of an oxidised guanine does not change the oligomeric hSSB1 binding mechanism

We have previously established that oxidised hSSB1 (referred to as ‘oligomeric hSSB1’ in this study; a mixture of monomers, dimers, tetramers and larger oligomers of hSSB1 protein based on our previous work ^21^) is required in cells to repair damaged DNA containing an 8-oxoG lesion ^19^. To initially test if the presence of an 8-oxoG base changes the molecular details of protein binding, HSQC experiments of oligomeric ^15^N-hSSB1_1-123_ were recorded in the presence (oligo(dT)_5_oxoG) with the sequence of [5′-TToxoGTTT-3′]) and absence (oligo(dT)_6_) of 8-oxoG, respectively (Figure 1A, B). Both the direction and the magnitude of the shift in the backbone moieties are conserved, suggesting that the presence of the oxidised guanine base does not affect the underlying molecular mechanism. This result is not unexpected, as DNA recognition by hSSB1 is modulated by base-stacking with ssDNA bases ^20^ and the oxidation of the guanine base does not alter its ability to stack with any of the key aromatic residues in hSSB1.

**Figure 1.**
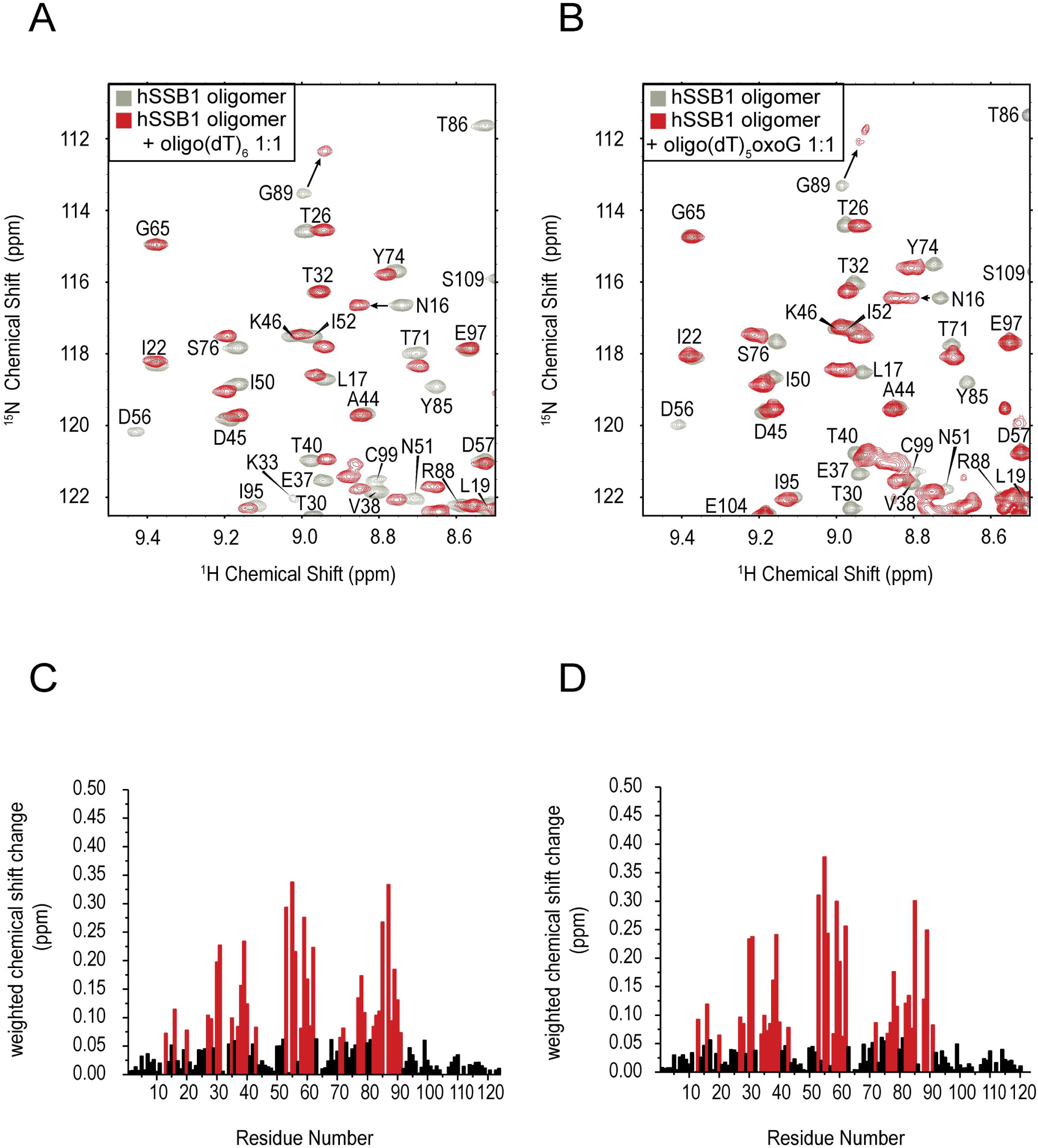
Oligomeric hSSB1 binding to 8-oxoG containing ssDNA. **A-B**. Section of ^15^N HSQC spectra of oligomeric hSSB1_1-123_ in the absence (grey) and presence (1:1; red) of oligo(dT)_6_ ssDNA (5′-TTTTTT-3′) (A) and 5ToxoG ssDNA (5′-TT(8-oxoG)TTT-3′) (B), respectively. **C-D**. Weighted backbone chemical shift changes of HN and N ^22^ atoms for hSSB1_1-123_ upon binding to oligo(dT)_6_ (C) and 5ToxoG (D) ssDNA, respectively. Residues that exhibit higher than average chemical shift changes are highlighted in red. Note the similarity of the chemical shift profiles of panels C and D.

### Oligomeric hSSB1 exhibits different binding kinetics compared to monomeric protein

The oxidation of hSSB1 is an essential prerequisite for oxidative damage repair and mutations that inhibit hSSB1 oligomer formation prevent the removal of damaged bases as previously shown ^16,19^. To address the question as to why hSSB1 needs to be oxidised in order to mediate the BER process, we next sought to determine the differences in ssDNA binding between oligomeric and monomeric hSSB1 by measuring the interaction kinetics using surface plasmon resonance (SPR) measurements. As seen in Figure 2, oligomeric hSSB1_1-123_ recognises oligo(dT)_12_ ssDNA with significantly different binding kinetics (in particular in relation to the off-rate) and a ∼7-fold decrease in the dissociation constant compared to the monomeric protein. Given local concentrations of hSSB1 can vary significantly in a cellular environment, this increase in binding affinity might explain why oligomeric and not monomeric proteins are required to facilitate oxidative damage repair. These data also provide strong evidence that ssDNA binding of oligomeric hSSB1 takes place by direct binding of hSSB1 oligomers (e.g., dimers or tetramers) as opposed to successive addition of dissociated monomers.

**Figure 2.**
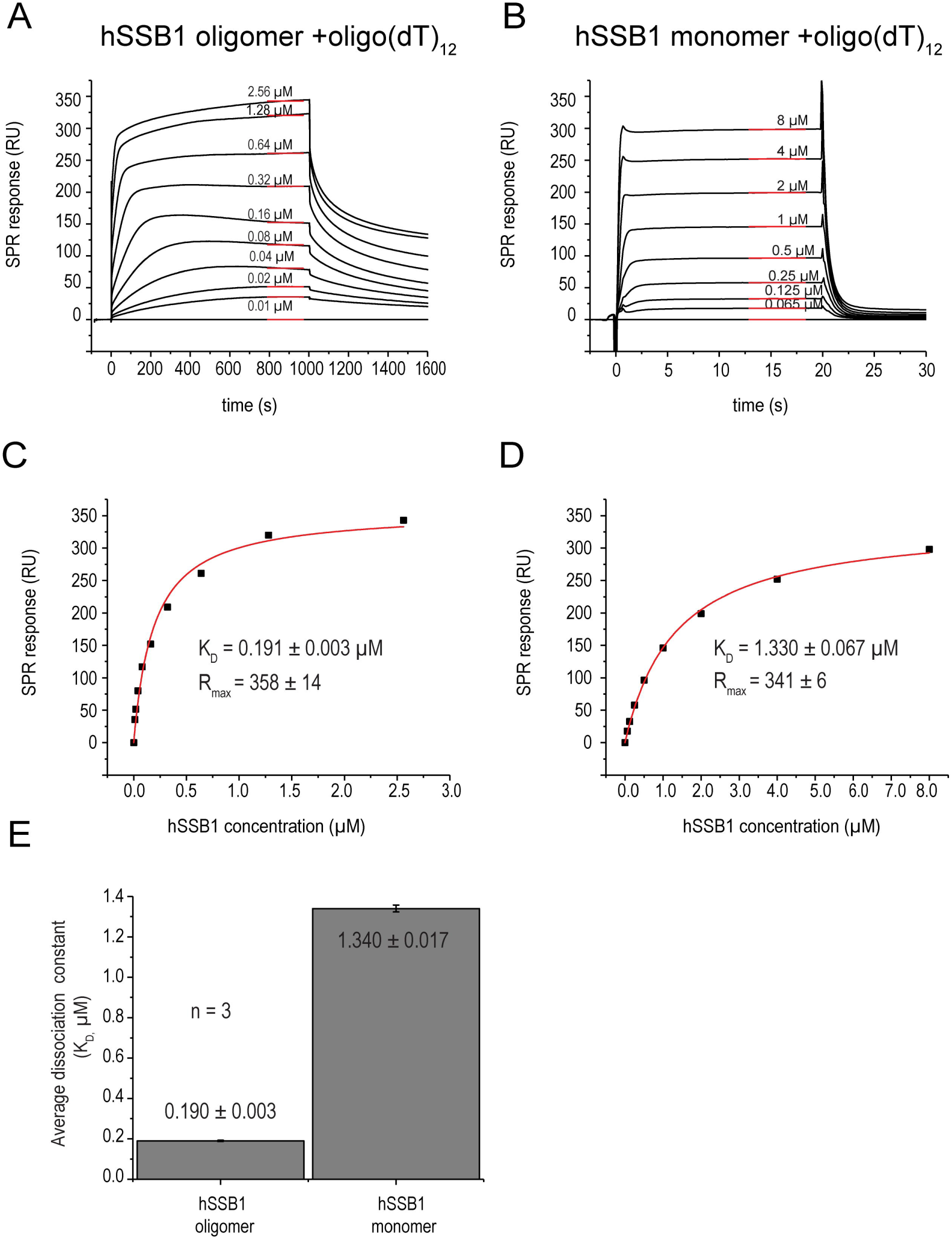
SPR analysis of binding of oligomeric hSSB1 to oligo(dT)_12_ ssDNA. Representative SPR binding curves of interaction between oligomeric (A) and monomeric hSSB1_1-123_ (B), respectively (concentration relates to monomers; note the different timescales used for data collection), with oligo(dT)_12_ ssDNA. **C-D**. Equilibrium values (as indicated by the red lines in panels A and B) for both oligomeric and monomeric proteins were plotted against the hSSB1_1-123_ concentration and fitted to the Langmuir equation (steady-state analysis). **E**. Dissociation constants were calculated from three independent experiments for both oligomeric and monomeric protein.

### The ssDNA binding interface is conserved between oligomeric and monomeric hSSB1

Next, to establish structurally how oligomeric hSSB1 proteins recognise ssDNA, we carried out HSQC experiments using ssDNA with varying lengths under reducing and oxidising conditions (Figure 3-5). Overall, protein residues that exhibit substantial chemical shift changes upon addition of oligo(dT)_6_, oligo(dT)_12_ and oligo(dT)_23_ ssDNA (Figures 3-5, panels A and B) are largely identical for both reduced and oxidised hSSB1. This is also indicative in the profiles of weighted chemical shift changes; residues undergoing significant changes are coloured in red in Figures 3-5 (panels C and D). Notably, all of the interacting residues are found in the known ssDNA binding surface for the short ssDNA (oligo(dT)_6_; see also Touma et al. ^20^) and no additional contacts between hSSB1 and ssDNA were observed in the binding complexes with the longer oligonucleotides (Figure 5 and 6). These data strongly suggest that even for longer ssDNA the interaction of oligomeric hSSB1 occurs via the established binding interface.

**Figure 3.**
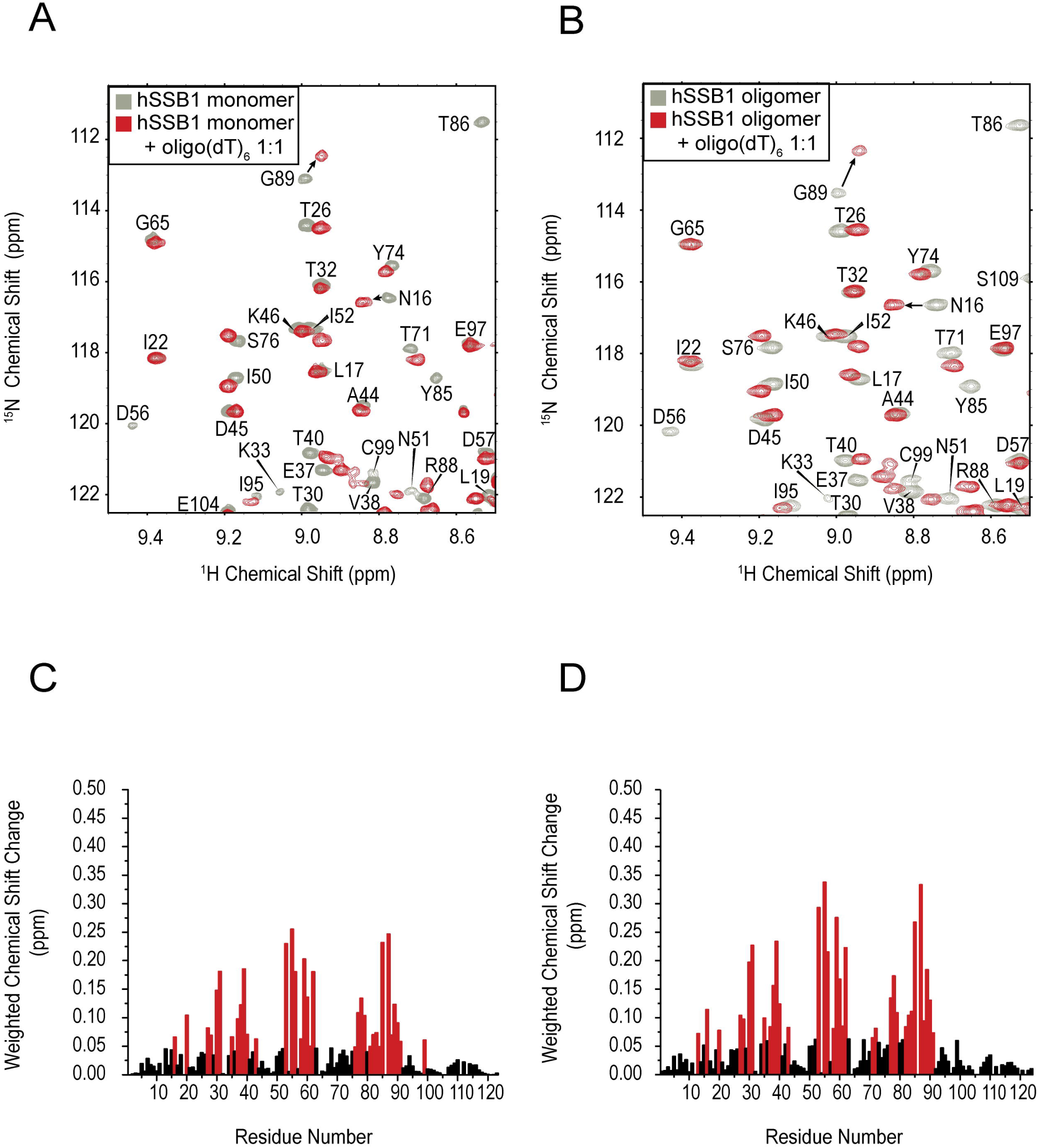
Difference of oligomeric versus monomeric hSSB1 binding to oligo(dT)_6_ ssDNA by 2D NMR HSQC experiments. Portion of ^15^N HSQC spectra in the absence and presence of oligo(dT)_6_ ssDNA (1:1) with monomeric (**A**) or oligomeric (**B**) hSSB1_1-123_, respectively. Difference in the weighted chemical shift changes between unbound and ssDNA bound hSSB1 monomer (**C**) and oligomer (**D**), respectively. Residues exhibiting changes that are significantly higher than average are highlighted as red bars.

**Figure 4.**
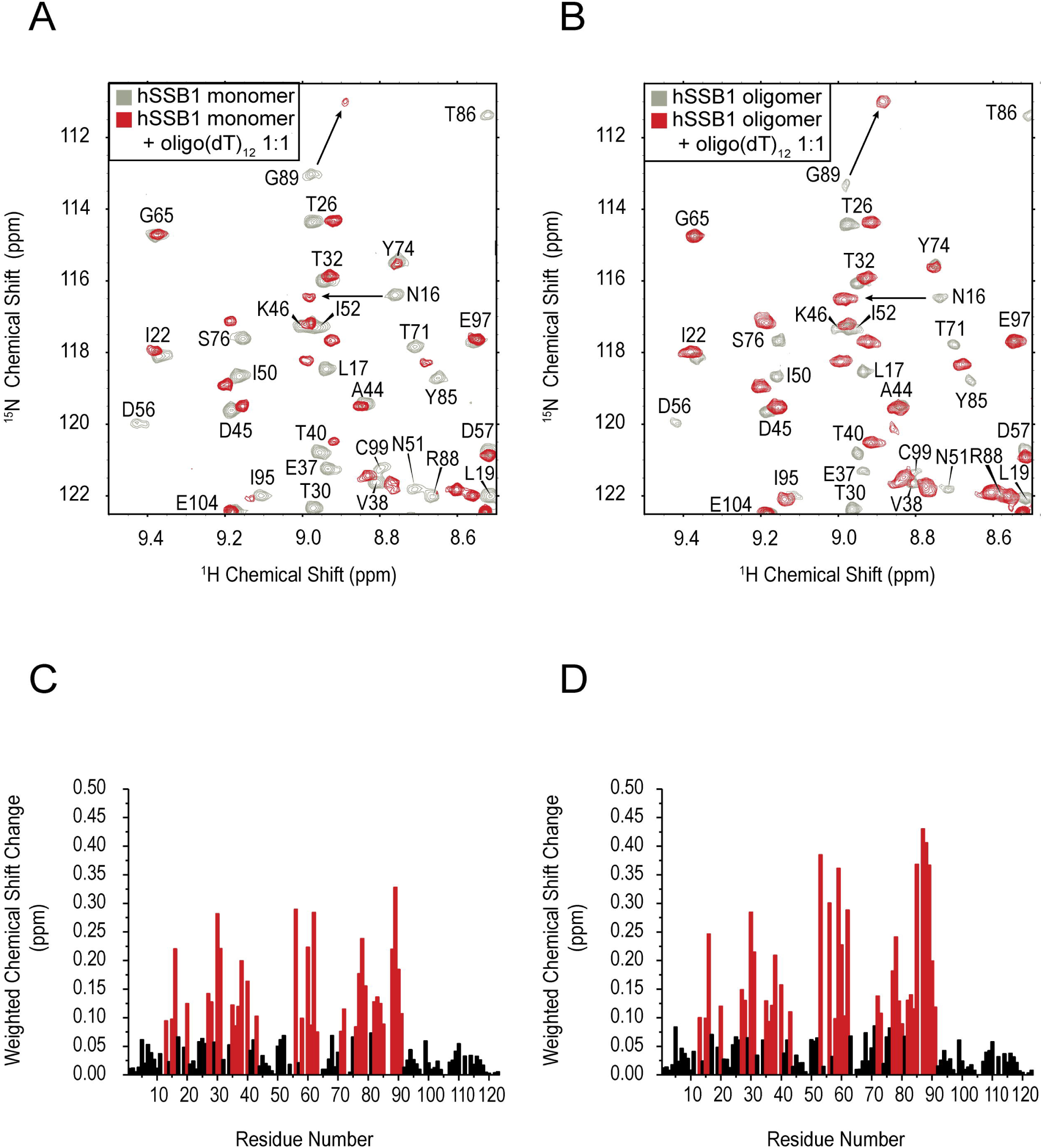
Difference of oligomeric versus monomeric hSSB1 binding to oligo(dT)_12_ ssDNA by 2D NMR HSQC experiments. Portion of ^15^N HSQC spectra in the absence and presence of oligo(dT)_12_ ssDNA (1:1) with monomeric (**A**) or oligomeric (**B**) hSSB1_1-123_, respectively. Difference in the weighted chemical shift changes between unbound and ssDNA bound hSSB1 monomer (**C**) and oligomer (**D**), respectively. Residues exhibiting changes that are significantly higher than average are highlighted as red bars.

**Figure 5.**
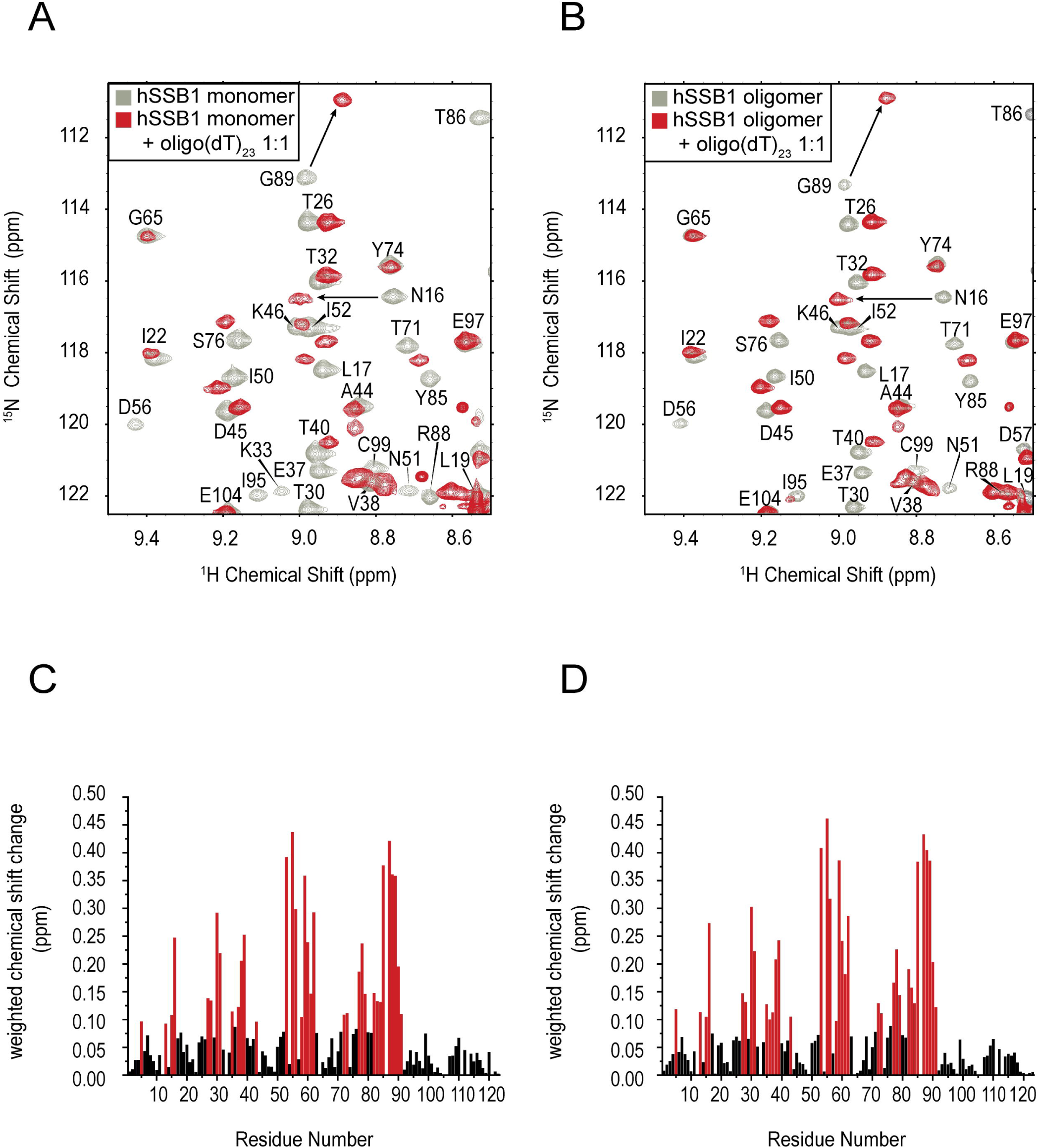
Difference of oligomeric versus monomeric hSSB1 binding to oligo(dT)_23_ ssDNA by 2D NMR HSQC experiments. Portion of ^15^N HSQC spectra in the absence and presence of oligo(dT)_23_ ssDNA (1:1) with monomeric (**A**) or oligomeric (**B**) hSSB1_1-123_, respectively. Difference in the weighted chemical shift changes between unbound and ssDNA bound hSSB1 monomer (**C**) and oligomer (**D**), respectively. Residues exhibiting changes that are significantly higher than average are highlighted as red bars.

**Figure 6.**
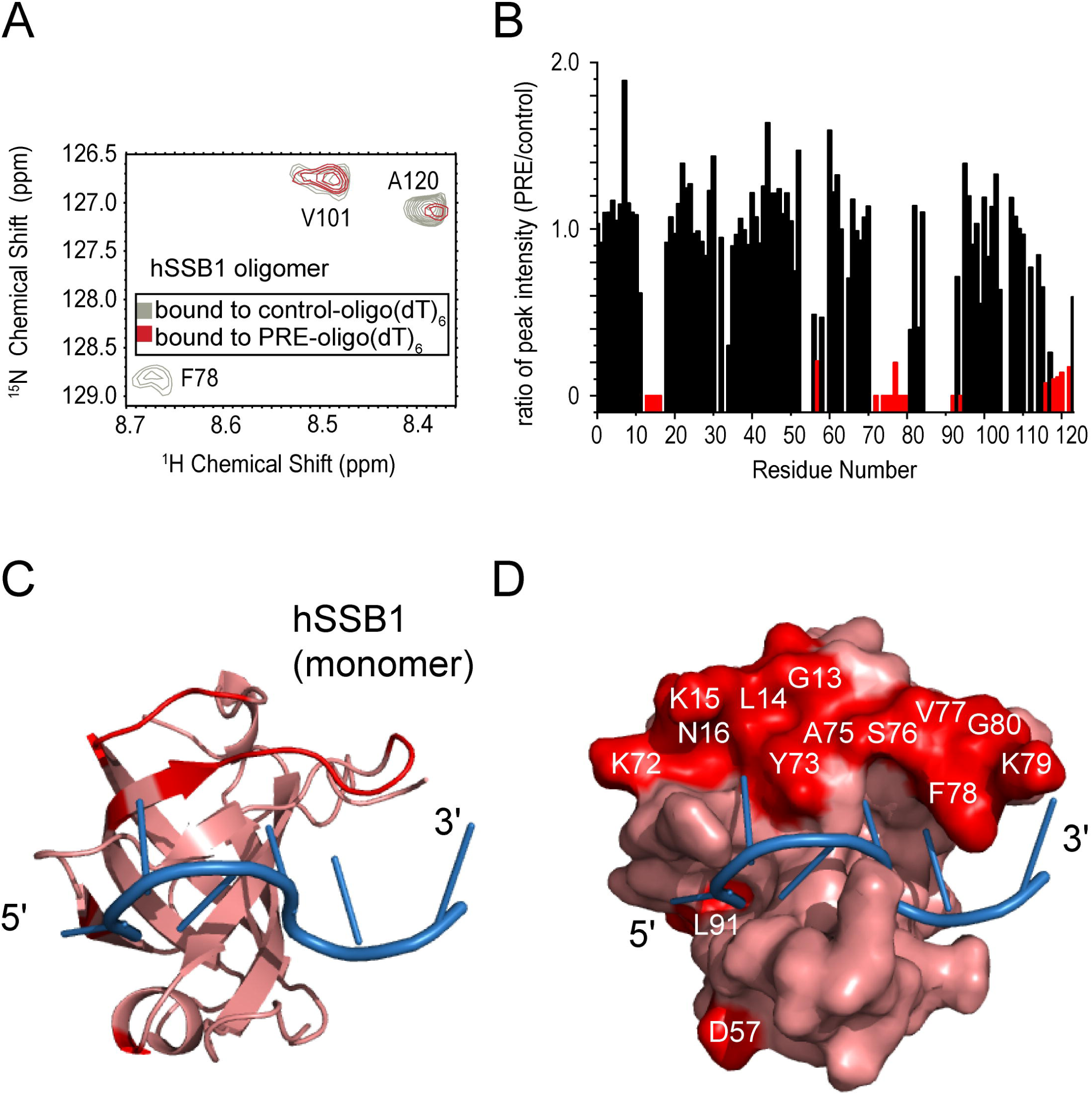
PRE analysis of oligomeric hSSB1 binding to ssDNA. **A**. Section of ^15^N HSQC spectra of a 1:1 mixture of hSSB1_1-123_ with paramagnetic nitroxide spin-labelled ssDNA [5′-G’(PRE)’GTTTTTT-3′] (PRE-oligo(dT)_6_; red) and diamagnetic non-labelled ssDNA [5′-G’GTTTTTT-3′] (control-oligo(dT)_6_; grey). Note that signals originating from residues that are close to the paramagnetic centre either disappear or are significantly reduced under paramagnetic conditions. **B**. Ratio of HSQC peak intensities of hSSB1_1-123_ residues under paramagnetic (PRE-oligo(dT)_6_) versus diamagnetic (control-oligo(dT)_6_) conditions. Ratios from signals of residues that exhibit a decrease significantly smaller (average minus two times standard deviation) than the average ratio or disappear entirely (ratio equals zero) are indicated in red in panel B. These residues (coloured in red) were mapped onto our published hSSB1-ssDNA model ^20^, visualised as cartoon or surface representation (**C-D**). Note that affected residues are located along the ssDNA binding site and close to both the 3′ and the 5′ end of the ssDNA.

### Oligomeric hSSB1 can bind ssDNA in both binding polarities

In previous work, we have shown that the OB domain of the SSB from *Sulfolobus solfataricus* (SsoSSB) interacts with ssDNA with a specific binding polarity ^23^. Both hSSB1 and SsoSSB use a combination of base-stacking between aromatic residues and ssDNA bases, as well as some hydrogen bonding to bind ssDNA. While base-stacking does not confer any directionality on ssDNA binding, most SSBs have a preferred binding orientation ^24^. Although both our NMR structure ^20^ and the published crystal structure ^18^ indicate a preferred binding polarity, no other experimental data have been obtained to exclude the possibility that a second conformer, which binds ssDNA in the reverse direction to that observed in the published structures, exists.

To address this, we carried out a semi-quantitative analysis using paramagnetic resonance enhancement (PRE) labelled ssDNA [5′-G’(PRE)’GTTTTTT-3′] (see experimental section for more details), in analogy to our published approach with SsoSSB ^23^ (Figure 6). Figure 6A depicts an overlay of a HSQC spectrum of oligomeric ^15^N-hSSB1_1-123_ in the presence of unlabelled ssDNA (diamagnetic conditions, labelled ‘control’ in Figure) onto a HSQC spectrum of ^15^N-hSSB1_1-123_ in the presence of PRE-labelled ssDNA (paramagnetic conditions, labelled ‘PRE’ in Figure). Calculation of the ratio between peak intensities of ‘PRE’ and ‘control’ conditions (Figure 6B) revealed residues that either exhibit no signal in the presence of PRE-ssDNA or undergo substantial decreases in their peak intensities (indicated as red bars in the Figure) and are thus located in close proximity to the PRE-label. These residues were mapped onto our published structural model of monomeric hSSB1 in complex with oligo(dT)_6_ ^21^ (coloured in red in Figure 6C, D). Notably, all residues that are within a small distance from the paramagnetic nitroxide spin-label are found in regions along the ssDNA binding site, with some of the affected residues located proximal to the 3′ end and others proximal to the 5′ end of the ssDNA binding surface (Figure 6C, D). Given the PRE-label is covalently bound to the 5′ end of the ssDNA, these data strongly indicate that, as opposed to SsoSSB, oligomeric hSSB1 is able to bind to ssDNA utilising both binding polarities.

## Discussion

### Comparison of ssDNA binding of oligomeric hSSB1 to *E. Coli* SSB

Our hSSB1 tetramer model ^21^ was initially based on comparisons with the tetramer formed by the OB domains from *E. Coli* (EcSSB) ^25^. Despite the structural similarities between the respective protein tetramers, the actual alignment when forming tetramers is significantly different between the two proteins. As opposed to the hSSB1 tetramer, the ssDNA binding grooves in the EcSSB tetramer (in the (EcSSB)_65_ binding mode) do not form a continuous binding surface, and to maintain the same binding polarity the strands of ssDNA wrap around the four monomers (Figure 7A; PDB ID 1EYG). In contrast, our NMR data (Figures 3-6) demonstrate that oligomeric hSSB1 can recognise ssDNA in both binding orientations using an overall binding interface that is different to EcSSB. In addition, the presence of the C99-C99 disulphide bond in the hSSB1 tetramer presents an obvious obstacle to ssDNA binding in this region and would prevent the ssDNA from interacting with the hSSB1 tetramer as in the conformation seen in EcSSB.

**Figure 7.**
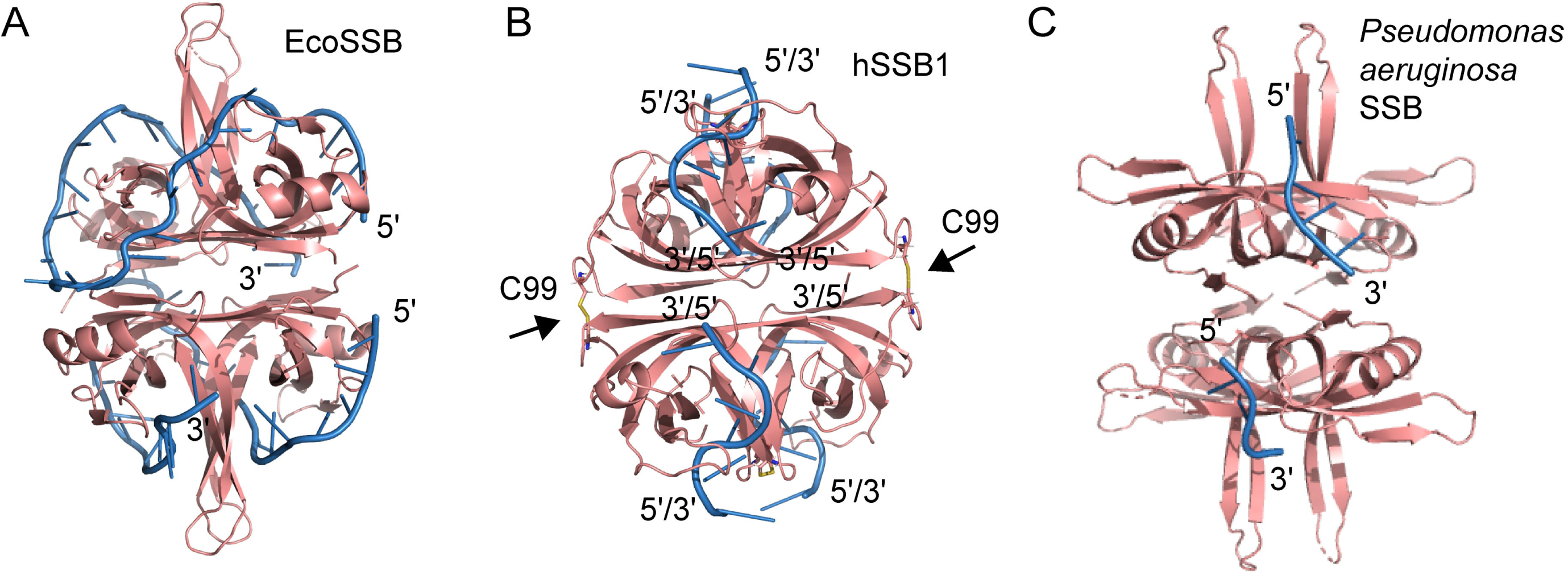
Comparison of ssDNA binding of hSSB1 tetramer to other SSBs. **A**. Crystal structure of the *E. Coli* SSB tetramer (EcSSB; salmon) bound to ssDNA (blue) (PDB ID 1EYG) ^33^. **B**. Structural model of the hSSB1 tetramer (salmon) taken from ^21^ with the oligo(dT)_6_ ssDNA (blue) overlaid from the ssDNA bound monomeric structure published in ^20^ with all possible ssDNA binding polarities indicated. The disulphide bond between C99 residues is indicated. Note the significant difference in ssDNA binding between EcSSB and hSSB1. **C**. X-ray crystal structure of tetrameric *P. aeruginosa* SSB (salmon) bound to ssDNA (blue) at 1.91 Å resolution (PDB ID 6IRQ) ^30^. Note that both ssDNA binding polarities exist in the structure (only the relevant ssDNA from PDB ID 6IRQ is shown).

We have previously also calculated a structural model of monomeric hSSB1 bound to ssDNA (oligo(dT)_6_) ^20^. To visualise where in the tetrameric structure ssDNA binding occurs based on our NMR data, our published model ^20^ was overlaid onto all four monomers within the tetramer model (Figure 7B); the ssDNA is shown in blue (the arrows indicate the position of the C99-C99 disulphide bond in the hSSB1 tetramer). In theory, every hSSB1 monomer can bind ssDNA in both binding orientations as indicated in Figure 7B, increasing the number of possible structural configurations.

### ssDNA binding polarity

The number and sequence of amino acids present in OB fold-containing proteins varies, however, there are several structural features that are common to all OB folds. OB folds from all three domains of life contain five anti-parallel β strands, coiled to form a β-barrel structure that is capped by an α helix between the third and fourth β strands at one end, and those that bind ssDNA contain a ssDNA binding cleft at the other ^26^. This ssDNA binding cleft is defined by two loops at the surface of the OB fold, perpendicular to the axis of the β-barrel ^24^.

Importantly, while the described ssDNA interaction interface is utilised throughout the entire OB domain family of proteins, the orientation that the DNA adopts (DNA binding polarity) is not conserved. However, the majority of OB fold proteins recognise ssDNA in what is defined by Theobald et al., ^24^ as the ‘standard binding polarity’ in which the DNA binds within the ssDNA binding cleft, with a polarity of 5′ to 3′ from strands β4 and β5 to strand β2 ^24^. One of two notable exceptions is the β subunit of *Oxytricha nova* telomere end binding protein (OnTEBP; PDB ID 1K8G) ^27^, which binds to ssDNA in the reverse orientation (‘reverse polarity’) ^24^. The second exception described by Theobald et al. ^24^ is EcSSB, bound to ssDNA in the (EcSSB)_65_ mode (PDB 1EYG).

Intriguingly, another stable homo-tetramer, the SSB from *Plasmodium falciparum* (PfSSB; PDB ID 3ULP) binds ssDNA adopting a topology that is highly similar to (EcSSB)_65_, however it does so using the standard polarity ^28^. In contrast, a more recently solved tetrameric OB fold-containing protein structure from *Bacillus subtilis* SsbA (BsSsbA; PDB ID 6BHX) ^29^ revealed ssDNA binding in the reverse polarity in analogy to EcSSB. These discrepancies could at least partly be explained by crystallographic artefacts in that one ssDNA binding orientation is more stable and therefore crystallised. However, this does not exclude the possibility that structures containing ssDNA with the opposite, or even both binding polarities at the same time, exist *in vivo*. Indeed, the latter is the case in one recently published structure, the tetrameric SSB from *Pseudomonas aeruginosa* (PaSSB; PDB ID 6IRQ; ^30^). Like hSSB1 (Figure 7B), in this crystal structure both ssDNA binding directionalities are observed (Figure 7C).

Orientation of ssDNA binding plays an important functional role for some SSBs, as is the case for human RPA in the context of nucleotide excision repair, where binding polarity appears to be crucial for positioning of the excision repair nucleases XPG and ERCC1-XPF onto the ssDNA ^31^. However, this contrasts with hSSB1, which forms oligomers as part of the response to oxidative damage. The main function of hSSB1 tetramers is to bind to oxoG lesions that exist in double-stranded DNA, a process that is then followed by the recruitment of hOGG1 that removes the lesion ^16,19^.

### Interactions with INTS3

We have previously shown that hSSB1 tetramer formation does not inhibit interaction with Integrator Complex Subunit 3 (INTS3), as two distinct interaction surfaces are utilised for binding ^21^. Our NMR data in this study reveal that ssDNA binding to oligomeric hSSB1 is mediated by protein residues that do not form part of the INTS3 recognition surface, thus theoretically enabling hSSB1 tetramer binding of INTS3 and ssDNA at the same time.

Interestingly, our structural data differs from a model described in a recently deposited preprint (Figure 6 in Jia et al. ^32^) incorporating hSSB1, ssDNA, INT3, and INTS6. In contrast to our and published data ^18^, this model ^32^ reveals direct contacts between the N-terminus of INTS3 and ssDNA. Further structural and biophysical studies will be required to shed light on the exact composition and structural details of the entire SOSS1 complex, comprising hSSB1 (SOSSB1), ssDNA, INTS3 (SOSSA), C9orf80 (SOSSC) and INTS6.

## Acknowledgements

We would like to thank Dr Ann Kwan for expert maintenance of the NMR spectrometers and Prof Nicholas Dixon for critical reading of the manuscript.

